# Human papillomavirus 16 E6 and E7 synergistically repress innate immune gene transcription

**DOI:** 10.1101/841007

**Authors:** Claire D. James, Christian T. Fontan, Raymonde Otoa, Dipon Das, Apurva T. Prabhakar, Xu Wang, Molly L. Bristol, Iain M. Morgan

## Abstract

Human papillomaviruses are causative agents in 5% of all cancers, including the majority of anogenital and oropharyngeal cancers. Downregulation of innate immune genes (IIGs) by HPV to promote the viral life cycle is well documented; E6 and E7 are known repressors of these genes. More recently we demonstrated that E2 could also repress IIGs. These studies have been carried out in cells over-expressing the viral proteins and to further investigate the role of individual viral proteins in this repression we introduced stop codons into E6 and/or E7 in the entire HPV16 genome and generated N/Tert-1 cells stably maintaining the HPV16 genomes. We demonstrate that E6 or E7 individually are not sufficient to repress IIG expression in the context of the entire HPV16 genome, both are required for a synergistic repression. The DNA damage response (DDR) is activated by HPV16 irrespective of E6 and E7 expression, presumably due to viral replication; E1 is a known activator of the DDR. In addition, replication stress was apparent in the HPV16 positive cells lacking E6 and E7, manifested by attenuated cellular growth and activation of replication stress genes. These studies lead us to the following model. Viral replication *per se* can activate the DDR following infection, and this activation is a known inducer of IIG expression which could induce cellular senescence. To combat this, E6 and E7 synergistically combine to manipulate the DDR and actively repress innate immune gene expression promoting cellular growth; neither protein by itself is able to do this.

**Importance:** The role of HPV16 in human cancers is well established; however, to date there are no anti-viral therapeutics that are available for combatting these cancers. To identify such targets, we must enhance understanding of the viral life cycle. Innate immune genes (IIGs) are repressed by HPV16, and we have reported that this repression persists through to cancer. Reversal of this repression would boost the immune response to HPV16 positive tumors, an area that is becoming more important given the advances in immunotherapy. This report demonstrates that E6 and E7 synergistically repress IIG expression in the context of the entire HPV16 genome. Removal of either protein activates the expression of IIGs by HPV16. Therefore, gaining a precise understanding of how the viral oncogenes repress IIG expression represents an opportunity to reverse this repression and boost the immune response to HPV16 infections for therapeutic gain.

## Introduction

HPV is the most common sexually-transmitted infection in the United States, with an estimated 80% of sexually active adults acquiring an HPV infection in their lifetime (1). Of the high-risk HPVs known to be causative in the development of cancer, HPV16 is the most prevalent genotype (2). HPV16 is causative in around 50% of cervical cancers and nearly 90% of HPV-related head and neck squamous cell carcinomas (HPV+HNSCCs). Further understanding of HPV16 and its life cycle is needed in order to develop novel anti-viral therapies targeting HPV16.

Much has been discovered relating to the viral oncoproteins E6 and E7 driving cellular growth and promoting infection (3–9). Most well-known are the targeting of cellular p53 and pRb by E6 and E7, respectively. E6 facilitates the degradation of p53 through its association with E6AP, a component of the ubiquitin-degradation pathway, whereas E7 binds to and disrupts the functions of pRb. The viral life cycle is dependent upon the differentiation program of epithelia; together E6 and E7 uncouple the process of differentiation and exit from the cell cycle to allow the virus to replicate and generate progeny virus.

In addition to targeting cellular proteins, both E6 and E7 can regulate transcription from the host genome of infected cells (6, 7, 10–12). Among the genes targeted for repression by E6 and E7 are the innate immune genes (11–16). Both viral oncoproteins have been shown to target expression of these genes, and such repression would promote viral infection by an overall suppression of the host immune response. In these studies, E6 and E7 have been over expressed from heterologous promoters. HPV16 infected cells also have an active DNA damage response (DDR) turned on at all times; E6, E7 and E1 have all been shown to activate this pathway (17–27). This provides another challenge the virus must overcome, as activation of the DDR stimulates the innate immune response (28). The DDR-mediated activation of the interferon response can occur in the absence of detectable foreign nucleic acids, such as following treatment of cells with etoposide. (29). Recently we demonstrated that repression of IIGs by HPV16 persists in HPV16 positive tumors and that this repression could contribute to the immune evasion by the HPV16 tumor (30, 31). We also recently demonstrated that, as well as E6 and E7, E2 can also repress expression of IIGs (30). Given that all of the IIG studies with E6, E7 and E2 have been done with over-expression of the viral proteins from heterologous promoters, we sought to investigate the contribution of each viral protein to IIG repression in the context of the entire HPV16 genome.

In order to investigate the contribution of the individual viral proteins to IIG repression, we generated mutant HPV16 genomes that had stop codons in E6 or E7, or both (see materials and methods). These were transfected into the hTERT immortalized foreskin keratinocyte cell line N/Tert-1 to establish stable cell lines containing wild type HPV16 and the assorted E6/E7 mutants. We have demonstrated that HPV16 represses IIGs in these cells (30, 31). These experiments required the use of already immortalized keratinocytes as HPV16 cannot immortalize primary keratinocytes in the absence of E6 and E7 expression. The results demonstrate that in the absence of either E6 or E7 there is an activation of IIG expression, not a repression. Removal of expression of both oncoproteins resulted in an additive activation of IIG expression. Strikingly, there is a strong synergism between E6 and E7 to repress IIGs as both proteins together can repress IIG expression in the context of cells containing the entire HPV16 genome. As HPV16 is known to activate the DDR, and an active DDR can activate expression of IIGs (28), we investigated the DDR in the wild type and mutant HPV16 genomes. We observed that the DDR is active in all HPV16 containing cells demonstrating that neither E6 nor E7 is required for activation of the DDR by HPV16. This activation likely results from the expression of E1 which is a known activator of the DDR (23, 25–27). We cannot generate stable cell lines over-expressing E1 by itself as it is toxic to cells. In addition, the cells lacking both E6 and E7 expression had severely attenuated cellular growth. All cells had activation of genes involved in managing replication stress. The results demonstrate that E6 and E7 synergize to repress expression of IIGs. They also demonstrate that activation of the DDR and induction of replication stress can occur in HPV16 cells independently of E6 and E7 expression, probably due to the replication factor E1.

## Results

### Establishment and characterization of mutant HPV16 genome N/Tert-1 cells

We introduced stop codons for E6 (residue C 110 mutated to a T) or E7 (residue T 584 mutated to an A) into the HPV16 genome, and both together. We prepared stable pooled N/Tert-1 cell lines containing the wild type and mutant HPV16 genomes. These were named N/Tert-1+HPV16, N/Tert-1+HPV16 E6 STOP, N/Tert-1+HPV16 E7 STOP and N/Tert-1+HPV16 E6E7 STOP. The presence of HPV DNA in the cells was confirmed by PCR using E6, E2 and E5 primers (Fig. 1A). While there was less DNA present in the mutants, levels were within 2-fold of each other between the different cell lines. The detection of E6, E2 and E5 demonstrates an intact early region in these cells. We next investigated RNA expression from the viral genome in these cell lines using qRT-PCR (Fig. 1B). There was less RNA present in the N/Tert-1+HPV16 E6E7 STOP cells indicating that these viral oncoproteins may actually promote transcription from the HPV16 genomes. However, in all cell lines there was significant expression of the viral genes and again, this was across the entire early region as E6, E2 and E5 expression is detected.

**Figure 1:**
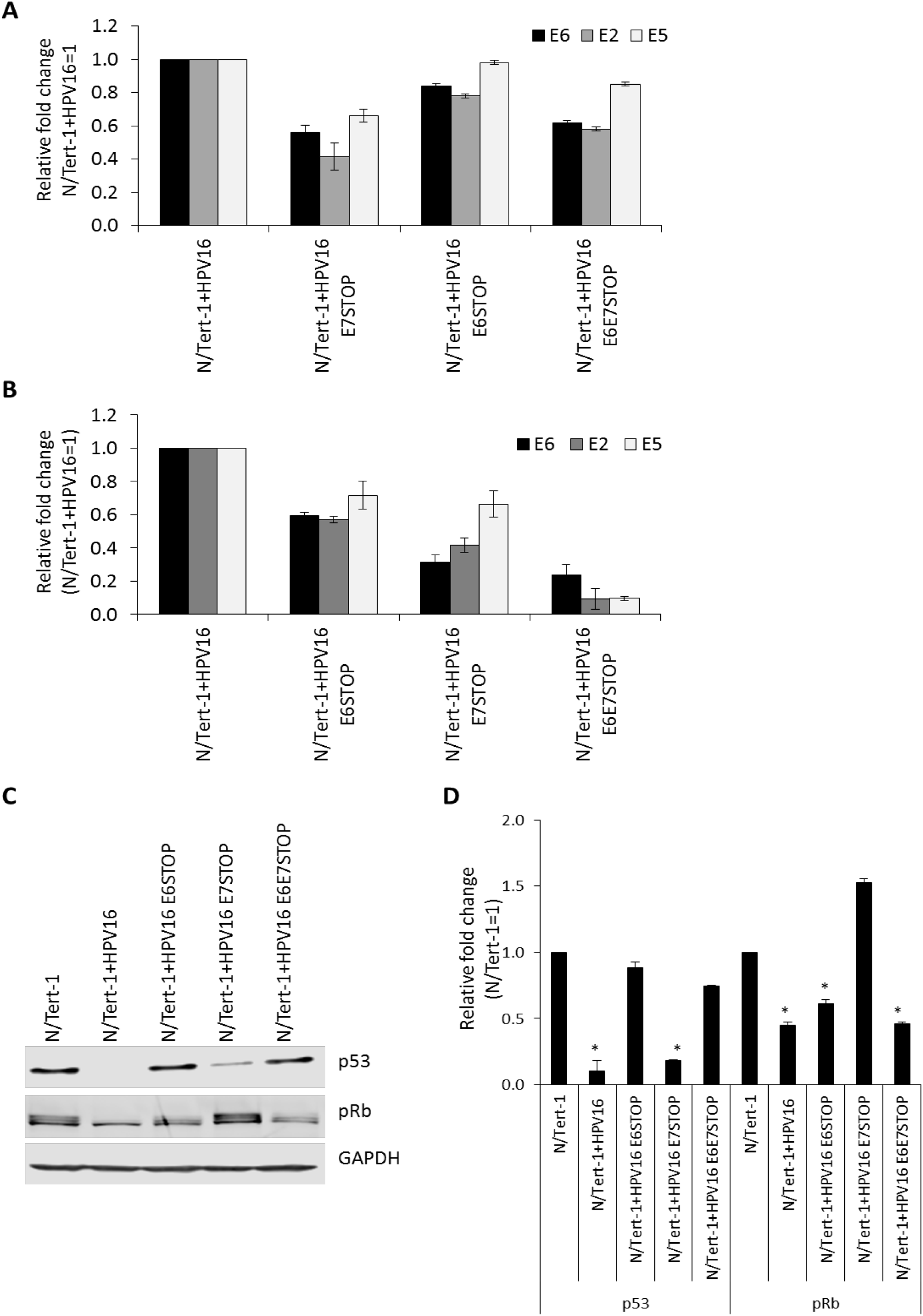
Initial characterization of mutant genome containing keratinocytes. A) qPCR analysis on DNA extracted from N/Tert-1+HPV16 cell lines. RNase-treated DNA was subject to SYBR green qPCR analysis, and ΔΔCt calculated using GAPDH housekeeping gene and normalized to N/Tert-1+HPV16. Error bars represent standard error of the mean of three individual experiments. B) qRT-PCR analysis for E2, E5 and E6 mRNA expression in N/Tert-1+HPV16, and mutant HPV16 genome containing cell lines. DNase treated RNA was subject to SYBR green qRT-PCR analysis and ΔΔCt calculated using GAPDH housekeeping gene and normalized to N/Tert-1+HPV16. Error bars represent standard error of the mean of three individual experiments. C) Western blot analysis for p53 and pRb in N/Tert-1, N/Tert-1+HPV16 and mutant HPV16 genome containing cell lines. GAPDH is shown as a loading control. p53 is downregulated in the presence of wild-type HPV16 and N/Tert-1+HPV16 E7STOP, but not in N/Tert-1+HPV16 E6STOP. pRb is downregulated in the presence of wild-type HPV16 and N/Tert-1+HPV16 E6STOP, but not in N/Tert-1+HPV16 E7STOP. Both pRb and p53 are unaffected in N/Tert-1+HPV16 E6E7STOP, compared to wild-type HPV16 genome containing cells. D) Western blots were visualized and quantified using Licor and calculated relative to parental N/Tert-1. Data represents the average of 3 independent experiments and error bars indicate standard error of the mean. * indicates p<0.05 compared to parental N/Tert-1 cells.

As markers for E6 and E7 activity, the expression levels of p53 and pRb were investigated (Fig. 1C). Both p53 and pRb are decreased in N/Tert-1+HPV16 cells and p53 levels are rescued to the same level as the parental N/Tert-1 cells in the E6 STOP and E6E7 STOP genomes. pRb is elevated when E7 is absent (E7 STOP), as would be expected. Surprisingly, pRb was down-regulated in N/Tert-1+HPV16 E6/E7 STOP cells. We confirmed that the E7 was mutated in these cells by DNA sequencing to eliminate the possibility of a plasmid mix up during transfection (not shown). These cells are stressed and have very slow growth (see below), therefore the presence of the replicating genome in the absence of E6 and E7 targets pRb for down-regulation by an as yet unknown mechanism. These blots were repeated and quantitated (Fig 1D).

### Absence of E6 and E7 expression increases innate immune gene expression in HPV16 containing cells

Our previous work in N/Tert-1 cells demonstrates that there is down regulation of innate immune gene expression at various stages of the interferon signaling pathway by HPV16 (30, 31). Following treatment of cells with interferon there is an activation of ISGF3 (interferon stimulated gene factor 3) which is a complex composed of STAT1, STAT2 and IRF9 (32–34). This transcription factor complex then enters the nucleus and binds to the control elements of interferon stimulated genes (ISGs) to activate their transcription. These ISGs are also downregulated in HPV16 positive N/Tert-1 cells (30, 31). These genes are repressed by E6, E7 and E2 and it was unclear what viral protein is responsible for this repression in the context of the entire HPV16 genome. Therefore, we investigated the expression of ISGs and STAT1-IRF9-STAT2 in the mutant genome HPV16 positive N/Tert-1 cells.

Fig. 2A demonstrates that the ISGs IFIT1, MX1 and OAS1 are repressed by HPV16 in N/Tert-1 cells, as we have reported (lanes 2, 7, 12). Strikingly, ablation of E6 expression (lanes 3, 8,13) or E7 expression (lanes 4, 9, 14) elevates ISG expression higher than that in control N/Tert-1 cells (lanes 1, 6, 11). In addition, ablation of both E6 and E7 expression (lanes 5, 10, 15) results in an additive increase in ISG expression. This regulation of RNA by the HPV16 genomes was reflected in the expression of the MX1 and IFIT1 proteins (Fig. 2B). This western blot was repeated and the results quantitated (Fig. 2C). These results demonstrate that both the RNA and protein levels of ISGs are elevated in HPV16 containing N/Tert-1 cells that have abrogated E6 and E7 expression, when compared with parental control cells.

**Figure 2:**
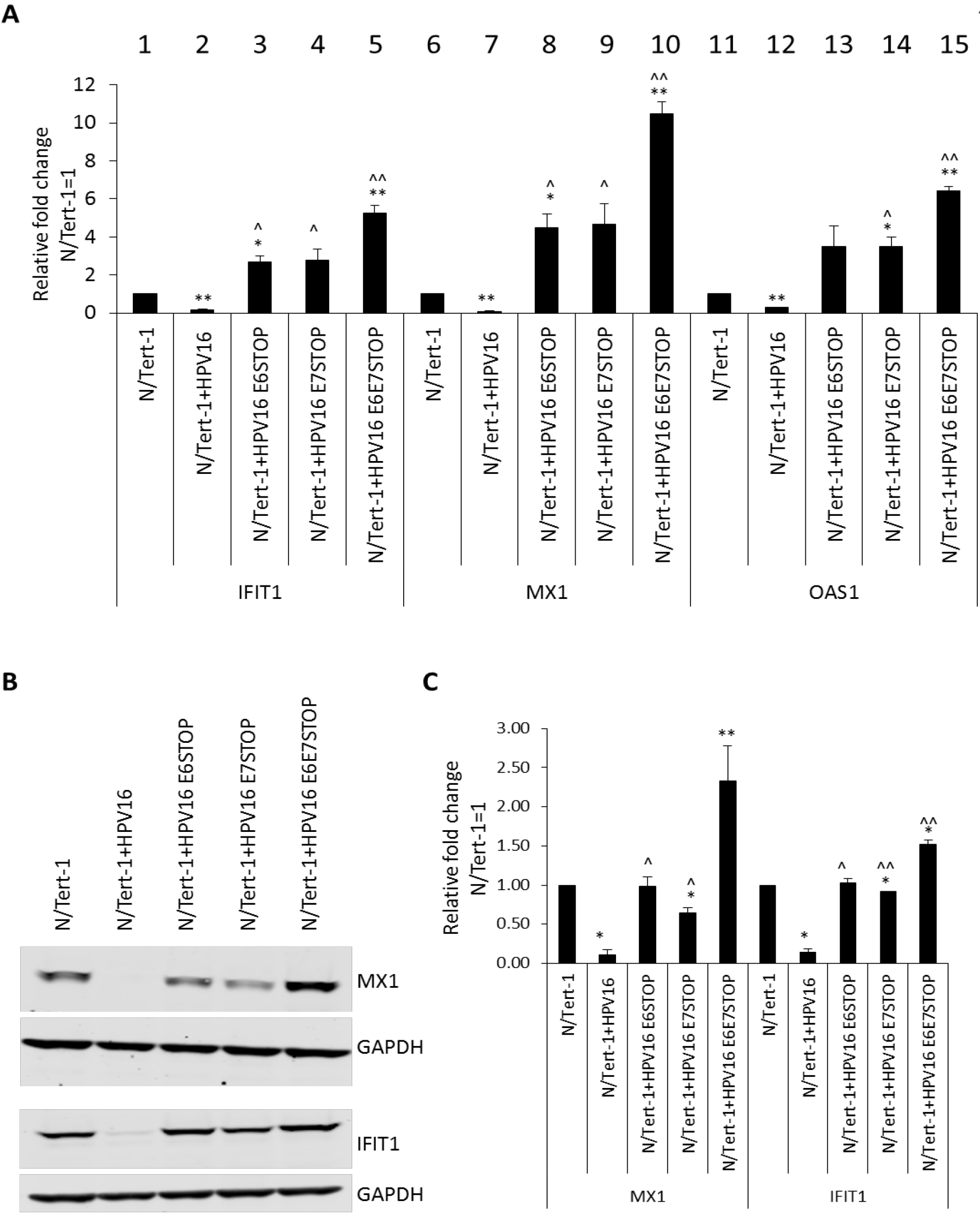
Absence of E6 or E7 expression increases innate immune gene expression in HPV16 containing cells. A) Quantification, by qRTPCR, of interferon stimulated genes expression, IFIT1, MX1 and OAS1, in N/Tert-1, N/Tert-1+HPV16 and mutant genome containing N/Tert-1 cell lines. ΔΔCt calculated using GAPDH housekeeping gene and normalized to parental N/Tert-1.. Data represents the average of 3 independent experiments and error bars indicate standard error of the mean. *=p<0.05 and **=p<0.001 relative to N/Tert-1, ^=p<0.05 relative to N/Tert-1+HPV16 and ^^ p<0.001 relative to N/Tert-1+HPV16. B) IFIT1 and MX1 protein expression was observed by western blot in N/Tert-1 cell lines. Representative images of three individual experiments, GAPDH included as loading control. C) Western blots were visualized and quantified using Licor and calculated relative to parental N/Tert-1. Data represents the average of 3 independent experiments and error bars indicate standard error of the mean. * indicates p<0.05 and **=p<0.001 relative to N/Tert-1, ^=p<0.05 relative to N/Tert-1+HPV16 and ^^ p<0.001 relative to N/Tert-1+HPV16.

We looked at the expression of the ISGF3 components in the mutant cells (Figure 3). As previously reported, STAT1 and IRF9 are repressed by HPV16 in N/Tert-1 cells, while STAT2 is not (compare lanes 2, 7, 12 with 1, 6, 11) (30, 31). Removal of E6 or E7 abrogates HPV16 repression of STAT1 or IRF9 in N/Tert-1 cells. STAT1 RNA levels were elevated in the absence of both E6 and E7 expression (compare lane 10 with 7). Protein expression levels of STAT1 and IRF9 are similarly regulated (Fig. 3B). Any repression of these proteins is lost following abrogation of E6 or E7 expression. The experiment in Fig. 3B was repeated and the western blots quantitated (Fig. 3C). Unlike the large increase in STAT1 RNA (Fig. 3A), there was no statistically significant difference in STAT1 protein levels in the absence of E6 and E7 when compared with abrogation of either protein expression by itself.

**Figure 3.**
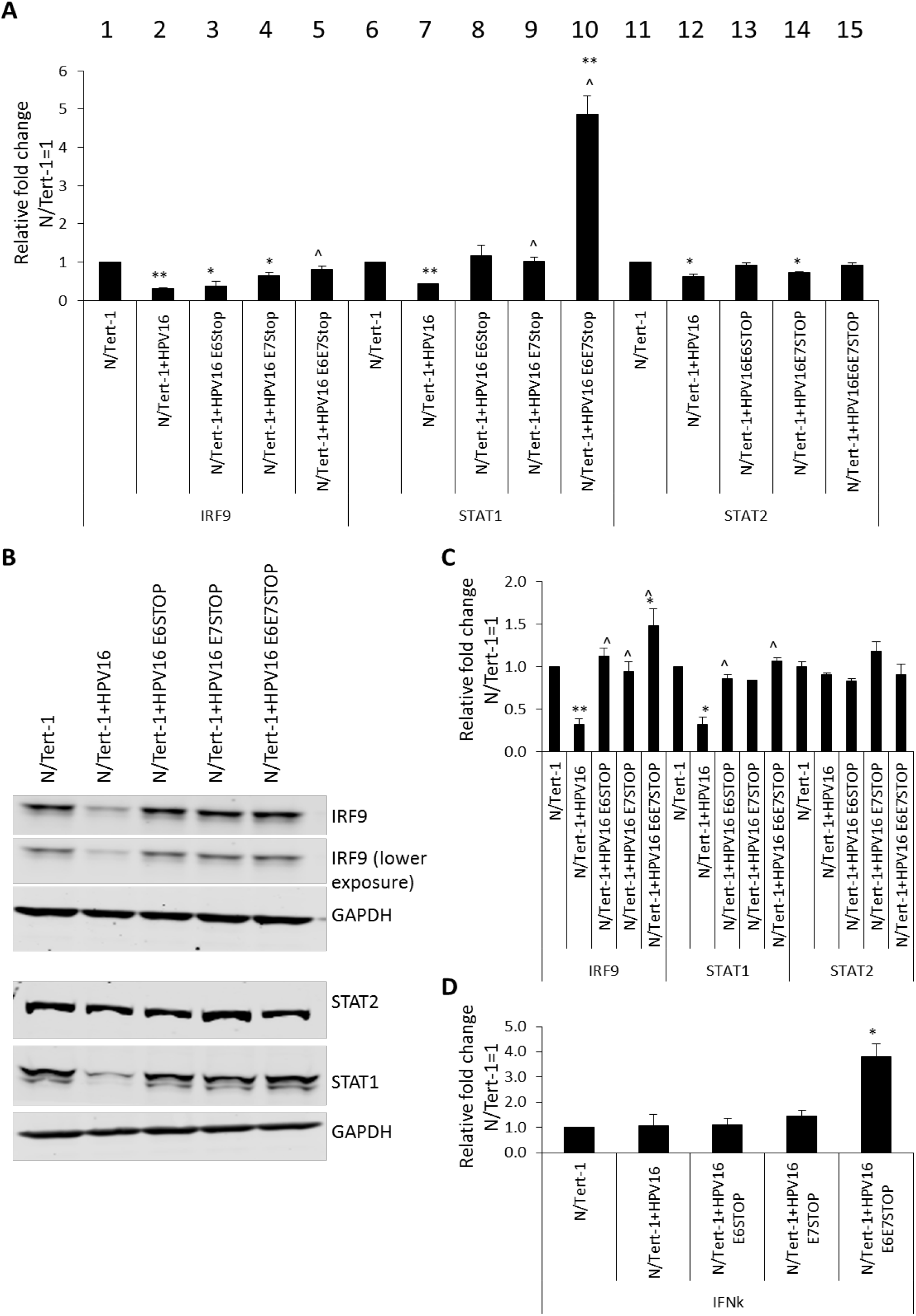
Absence of E6 and E7 expression affects upstream components of innate immune gene signaling in HPV16 containing cells. A) Quantification, by qRTPCR, of interferon stimulated genes expression, IRF9, STAT1 and STAT2 in N/Tert-1, N/Tert-1+HPV16 and mutant genome containing N/Tert-1 cell lines. ΔΔCt calculated using GAPDH housekeeping gene and normalized to parental N/Tert-1.. Data represents the average of 3 independent experiments and error bars indicate standard error of the mean.*=p<0.05 and **=p<0.001 relative to N/Tert-1, ^=p<0.05 relative to N/Tert-1+HPV16 and ^^ p<0.001 relative to N/Tert-1+HPV16. B) IRF9, STAT1 and STAT2 protein expression was observed by western blot in N/Tert-1 cell lines. Representative images of three individual experiments, GAPDH included as loading control. C) Western blots were visualized and quantified using Licor and calculated relative to parental N/Tert-1. Data represents the average of 3 independent experiments and error bars indicate standard error of the mean. * indicates p<0.05 and **=p<0.001 relative to N/Tert-1, ^=p>0.05 relative to N/Tert-1+HPV16 relative to N/Tert-1+HPV16. D) IFNκ expression quantified by qRTPCR. ΔΔCt calculated using GAPDH housekeeping gene and normalized to parental N/Tert-1. Data represents the average of 3 independent experiments and error bars indicate standard error of the mean. * indicates p<0.001 relative to N/Tert-1.

To determine whether the activation of STAT1 expression in the absence of E6 and E7 was due to an increase in interferon production we monitored the RNA levels for interferon genes. There was no detectable expression of interferon α or β in any of the N/Tert-1 cells. However, there was an increase in interferon κ RNA in the absence of both E6 and E7 expression when compared with parental cells (IFNκ, Fig. 3D). We previously observed a slight repression of IFNκ in HPV16 positive N/Tert-1 cells that we did not observe here, perhaps due to the use of clonal lines, rather than the pools used here. It is possible that the elevated IFNκ is responsible for the activation of STAT1 expression.

### The absence of functional E6 and E7 in HPV16 positive N/Tert-1 cells attenuates cellular growth

In doing the above experiments, we noticed that the N/Tert-1+HPV16 E6E7 STOP cells grew slower than the other N/Tert-1 cell lines. To quantitate this, we carried out a growth curve and demonstrated that this was indeed the case (Fig. 4). The presence of the HPV16 genome and the absence of E6/E7 expression resulted in dramatically attenuated growth when compared with parental N/Tert-1 cells. The N/Tert-1+HPV16 cells grew faster than parental N/Tert-1 cells as expected, and removal of E6 expression did not change this increased expression. The absence of E7 expression reduced the growth rate of the cells to that of the parental N/Tert-1 cells. However, the absence of both E6 and E7 synergized to dramatically reduce the growth of the N/Tert-1 cells.

**Figure 4.**
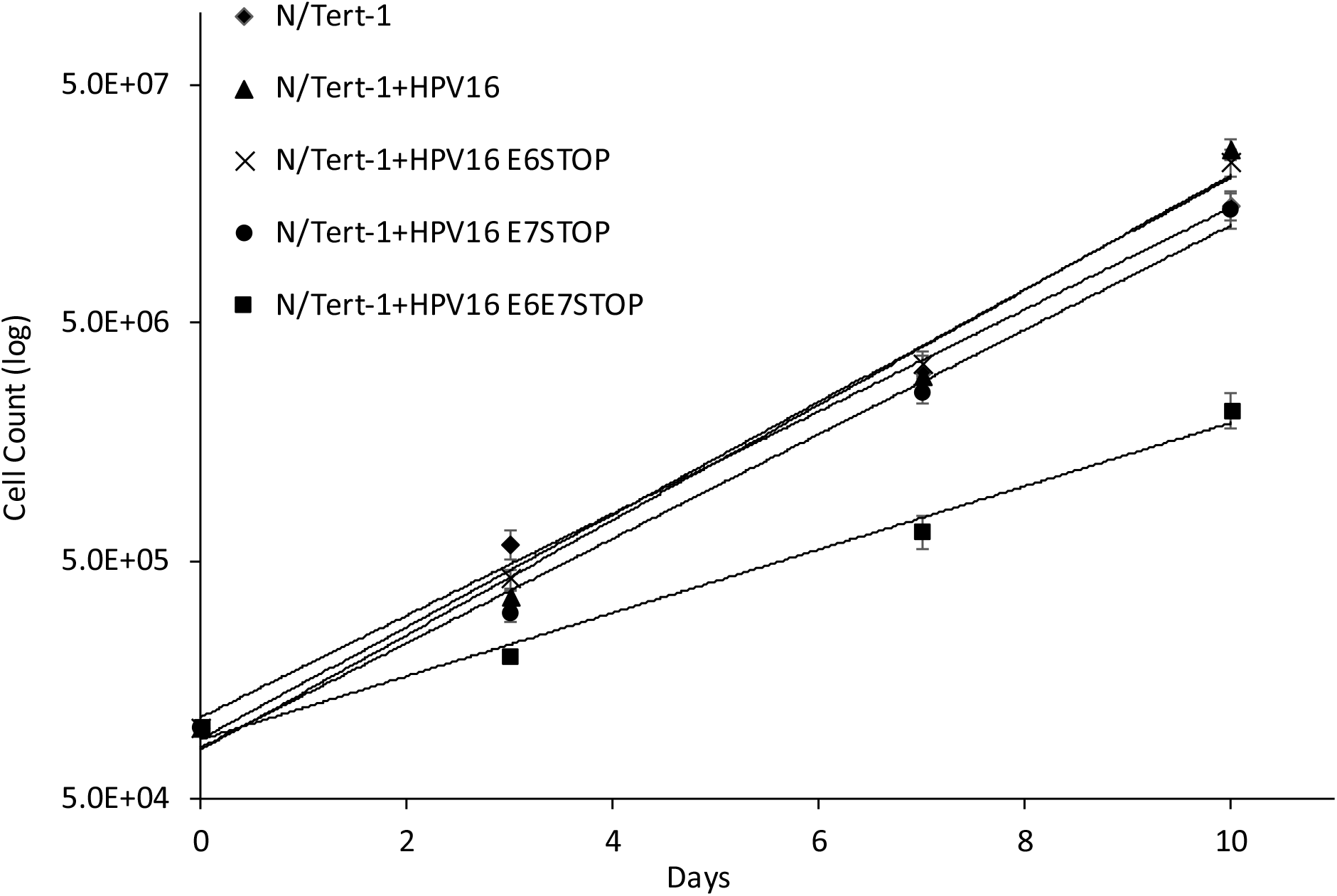
The absence of functional E6 and E7 in HPV16 positive N/Tert-1 cells attenuates cellular growth. To measure cell growth, cells were seeded in triplicate onto 10 cm dishes at a density of 3 × 10^5^ cells per dish, and grown to 80% confluency (typically 3 days). Cells were then harvested by trypsinization, stained with trypan blue and viable cells counted. 3 × 10^5^ cells per dish were replated and this was repeated 3 times in total. Data represents the average of 3 independent experiments and error bars indicate standard error of the mean. N/Tert-1+HPV16 grew significantly faster than N/Tert-1 and loss of E6 did not affect this. Removal of E7 resulted in the N/Tert-1 cells growing similarly to the parental N/Tert-1. Removal of E6 and E7 resulted in significantly slower growth of N/Tert-1 cells when compared with all other lines. In all analysis, p<0.05 to assign significance.

### The DNA damage response is activated in N/Tert-1 cells irrespective of E6/E7 expression

It is well established that HPV16 activates the DNA damage response (DDR) in keratinocytes, and that individual over-expression of E6 or E7 can also activate the DDR (17, 18, 20–22, 24, 35, 36). In addition, the E1 protein has also been shown to activate the DDR (23, 25–27). Given that the activation of the DDR pathway in non-HPV16 containing cells ordinarily results in growth attenuation, we investigated whether the DDR was activated in HPV16 cells in the absence of either E6 or E7 expression, or both. The levels of CHK1 and CHK2 RNA in N/Tert-1 cells are elevated by the HPV16 genome irrespective of E6/E7 status (Fig. 5A). This is reflected at the protein level (Fig. 5B). In addition, both CHK1 and CHK2 are activated in HPV16 positive N/Tert-1 cells irrespective of E6 and E7 status, as demonstrated by an increased activating phosphorylation of these proteins. Using immunofluorescence staining, we also observed that levels of γ-H2AX are increased in N/Tert-1 cells containing HPV16 irrespective of E6 and E7 levels (Fig. 5C).

**Figure 5:**
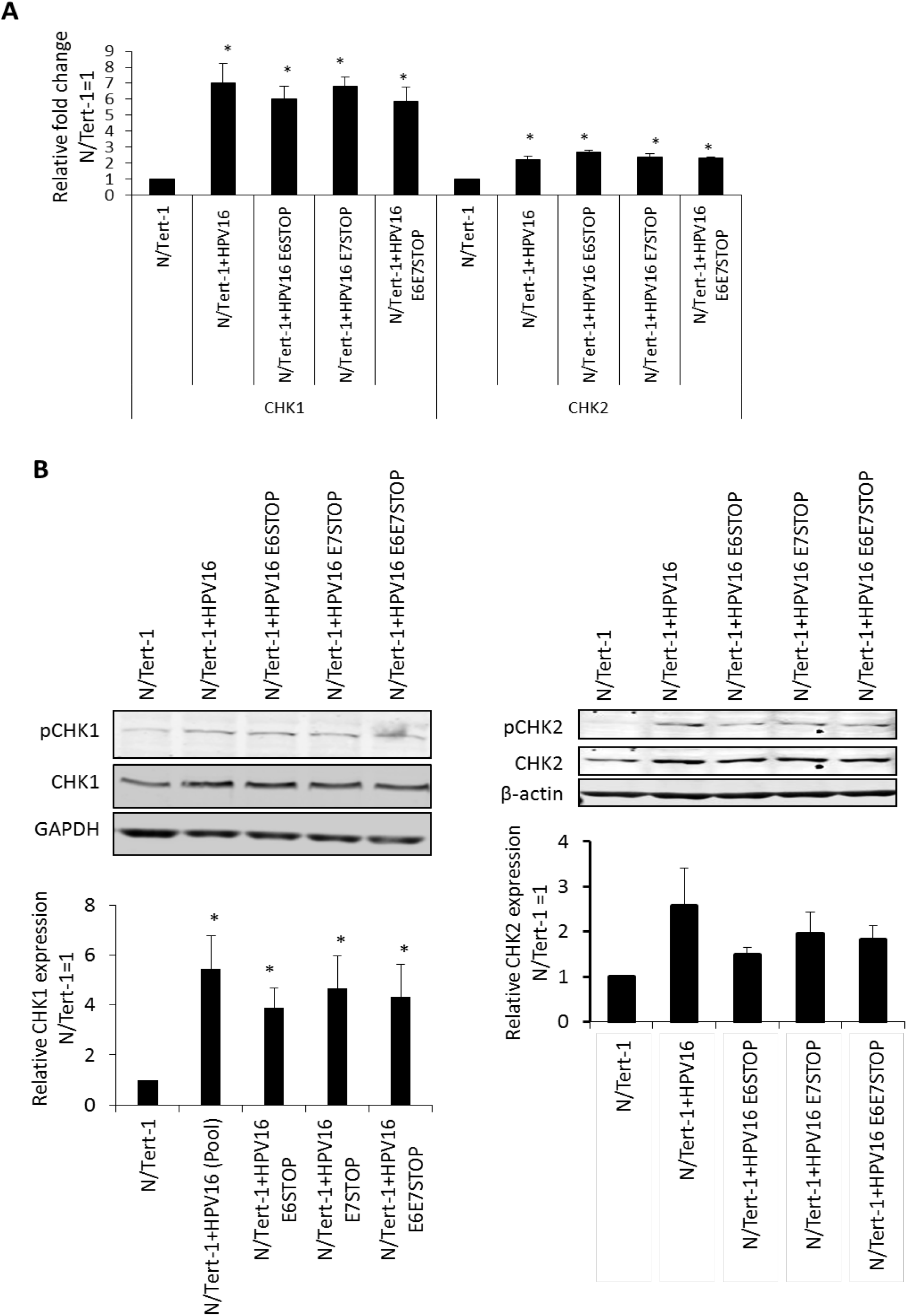
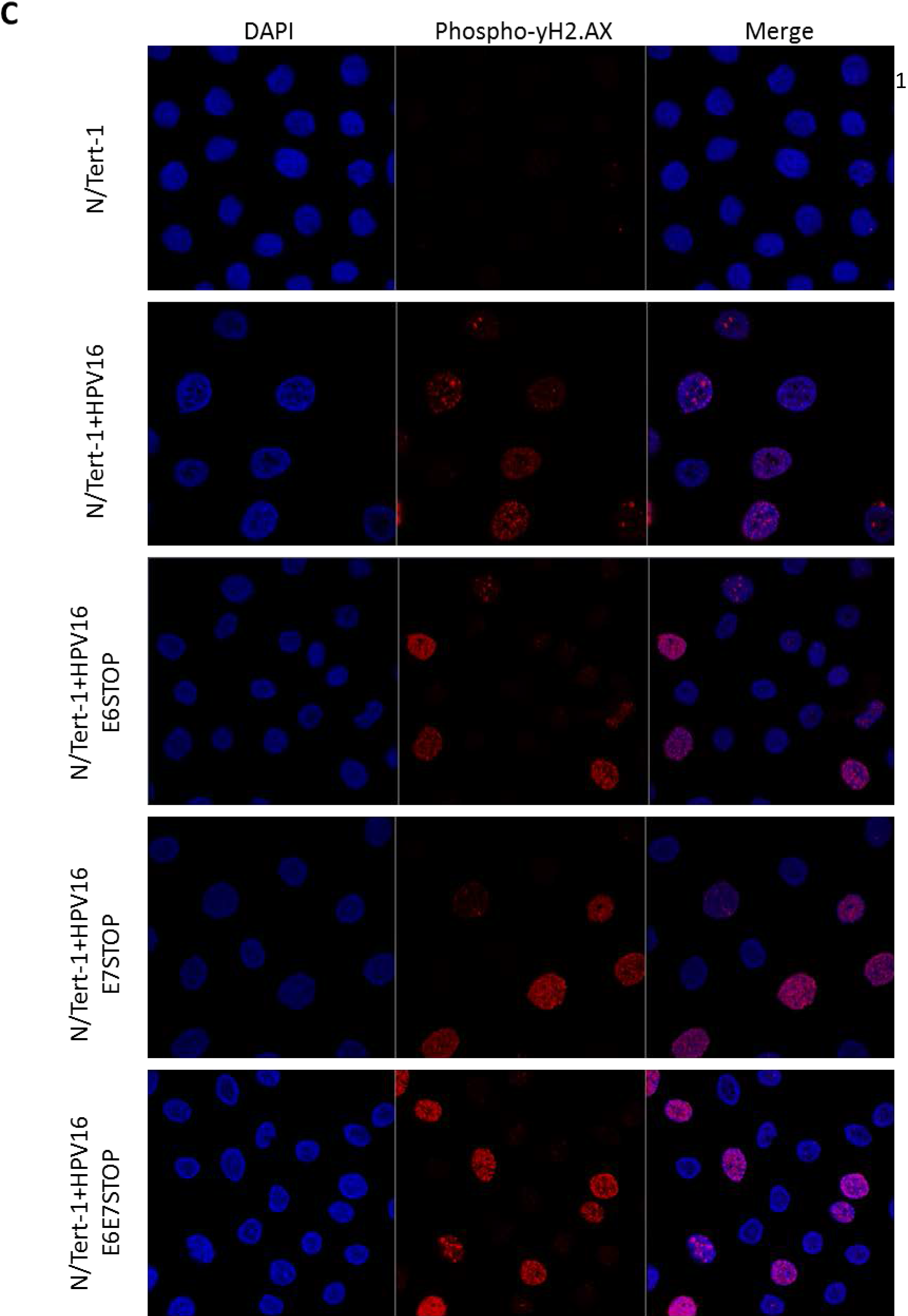
The DNA damage response is activated in N/Tert-1 cells irrespective of E6/E7 expression. A) qRTPCR analysis of CHK1 and CHK2 mRNA expression in N/Tert-1, N/Tert-1+HPV16, and mutant HPV16 genome containing cell lines. DNAse treated RNA was subject to SYBR green qRTPCR analysis and ΔΔCt calculated using GAPDH housekeeping gene and normalized to N/Tert-1. Error bars represent standard error of the mean of three individual experiments. Data represents the average of 2 or more independent experiments and error bars indicate standard error of the mean. * indicates p<0.05 relative to N/Tert-1. B) CHK1 and CHK2 protein expression and activation (observed as phosphorylated proteins) detected by western blot of N/Tert-1 cell lines. Representative images of three individual experiments, GAPDH included as loading control. Western blots were visualized and quantified using Licor and calculated relative to parental N/Tert-1. Data represents the average of 3 independent experiments and error bars indicate standard error of the mean. * indicates p<0.05 and **=p<0.001 relative to N/Tert-1, ^=p>0.05 relative to N/Tert-1+HPV16. C) Levels of DNA damage indicator phospho-γH2AX assessed by immunofluorescent staining of monolayer grown cells. Prior to staining, N/Tert-1 cell lines were grown to 70% confluency. Cellular DNA stained by inclusion of 4’,6-diamidino-2-phenylindole (DAPI). Images are representative of three individual experiments.

These results suggested that the N/Tert-1 cells containing HPV16 were under replication stress irrespective of E6/E7 status of the viral genome. We next investigated the RNA expression pattern of stress response genes in the N/Tert-1 cells containing the HPV16 genomes (Fig. 6). BRCA1, BRCA2 and DNA2 levels were all elevated in the HPV16 positive N/Tert-1 cells when compared with parental control, irrespective of E6 and E7 status. For ATR and Rad50 there were less dramatic increases in expression by any of the HPV16 genomes. The results investigating the DDR pathway in the N/Tert-1 cells containing the HPV16 genome demonstrates that replication stress and activation of the DDR is turned on by HPV16 independently from E6 and E7 expression.

**Figure 6:**
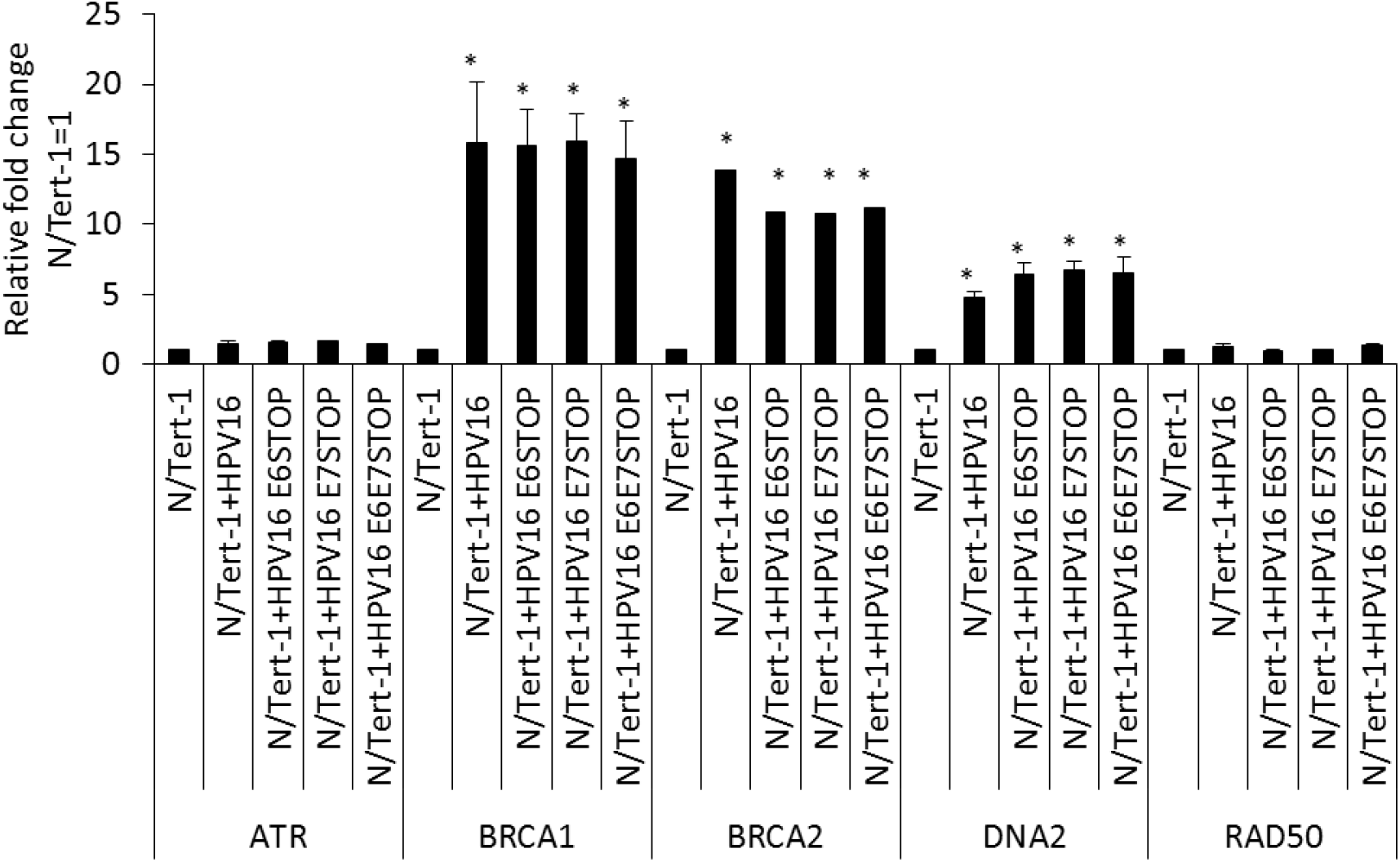
DNA Damage response gene expression is activated in N/Tert-1 cell lines containing HPV16 genomes. qRT-PCR analysis of DNA damage response genes in N/Tert-1, N/Tert-1+HPV16 and mutant HPV16 genome containing cell lines. DNAse treated RNA was subject to SYBR green qRTPCR analysis and ΔΔCt calculated using GAPDH housekeeping gene and normalized to N/Tert-1. Error bars represent standard error of the mean of three individual experiments. * indicates p<0.05 compared to parental N/Tert-1 cells.

## Discussion

Several labs have demonstrated repression of the innate immune response by high risk HPV in keratinocytes (13, 15, 16, 37). It is also established that, when E6 and E7 are overexpressed individually in keratinocytes, they can repress expression of innate immune genes (IIGs) (12, 38–40). Recently we demonstrated that HPV16 E2 can also repress IIG expression when overexpressed in keratinocytes (30). Given that all three viral proteins can repress innate immune gene expression when overexpressed, we sought to extend these studies to the HPV16 genome. To do this, we generated keratinocytes containing HPV16 with stop codons in either E6 or E7, or both together, to abrogate oncoprotein expression, and leaving the expression of E2 intact. Given that E6 and E7 are required for the generation of immortalized HPV16 containing primary cells we used foreskin keratinocytes immortalized by hTERT, N/Tert-1, for these experiments (41). Previously we have used these cells to demonstrate that E2 can regulate expression of host genes in keratinocytes, and that this regulation was relevant to the viral life cycle (30). The mutant and wild-type genome containing cell lines generated demonstrated the presence of HPV16 DNA with corresponding RNA expression from the viral genome. The N/Tert-1 cells with genomes containing stop codons in E6 or E7 demonstrated the predicted expression of the tumor suppressor proteins p53 and pRb; i.e. p53 is not degraded when E6 is non-functional, and pRb is expressed when E7 is non-functional. However, in the cells containing the genomes with stop codons in both E6 and E7 the levels of Rb are down regulated by an as yet unclear mechanism perhaps related to the replication stress these cells are under (Fig. 1).

We next investigated the expression of interferon stimulated genes (ISGs) in these cell lines and made several striking observations. First, HPV16 represses the expression of ISGs in N/Tert-1 cells, as we have previously reported (30, 31). Second, in the absence of either E6 or E7 there is an increase in the expression of the ISGs investigated to levels above what is observed in the vector control N/Tert-1 cells. Third, in the absence of both E6 and E7 there is an additive increase in the expression of the ISGs, increasing all of their expression levels to statistically significant higher levels than the parental N/Tert-1 cells. Fourth, this increase in RNA expression is reflected in protein expression; elevated levels of IFIT1 and MX1 in the absence of E6 and E7 was observed when compared with parental cells. Fifth, looking at the upstream ISGF-3 complex genes (the complex responsible for activating the ISGs) there is an enhanced increase in STAT1 levels when compared to N/Tert-1 cells where E6 and E7 are not expressed, although this is not reflected at the protein level. Sixth, there is an increased expression of IFNκ in the absence of both E6 and E7 when compared with all other N/Tert-1 cell lines.

These results demonstrate that, in the absence of E6 and E7, E2 cannot repress IIG expression alone. However, neither E6 nor E7 can repress these genes by themselves therefore there remains the possibility that E2 synergizes with E6 and E7 to repress IIGs in the context of the entire HPV16 genome. It is also possible that E2, E6 and E7 have roles to play in IIG repression during the different stages of the viral life cycle and this is currently under investigation.

It is clear that both E6 and E7 are required to suppress innate immune gene expression in the context of the entire HPV16 genome. While both proteins can repress genes when overexpressed individually, abrogation of expression of either results in a loss of innate immune gene repression (Figs. 2–3). This demonstrates a previously unidentified synergism between the two viral proteins in the context of the entire HPV16 genome. Both proteins have been shown to target the interferon signaling pathway using different mechanisms. E7 can bind to and inhibit the transactivating function of IRF1 (42), while E6 binds to IRF3 and inhibits its transcriptional activity (43) and can also interfere with JAK-STAT activation by binding Tyk2 (38). Our studies demonstrate that abrogation of both E6 and E7 expression results in an additive increase in the expression of ISGs. Therefore, the cross talk between E6 and E7 to suppress the innate immune response is synergistic, and the additive nature of ISG increase in the absence of these two proteins suggests independent pathways that are targeted by these viral proteins. The results presented here suggest that disruption of the mechanism that either of the viral proteins uses to suppress the innate immune response is a potential therapeutic target that would boost the host immune response against HPV infections.

The next question we addressed was what could be responsible for the elevated innate immune gene expression in the absence of E6/E7? The innate immune response is activated by detection of cytoplasmic DNA via the cGAS-STING pathway, cytoplasmic DNA can be detected in cells treated with DNA damaging agents (44). However, HPV16 E7 acts to combat activation of the innate immune response via the cGAS-STING pathway (45). A source for activation of the DDR in HPV16 cells lacking E6 and E7 is the E1 viral helicase that can activate the DDR by itself (23, 25, 26, 46). Although E6 and E7 can activate the DDR when overexpressed in cells, it is clear that in their absence HPV16 activates DDR pathways and we propose this is mediated by the viral replication factor E1 (Figs. 5&6). This activation of the DDR by HPV16 replication in the absence of E6 and E7 results in activation of the expression of innate immune response genes, perhaps via activation of the cGAS-STING pathway although we could not detect several markers for activation of this pathway in the N/Tert-1+HPV16 E6/E7 STOP cells (not shown). In support of the idea that there is increased aberrant DNA sensing in the absence of E6 and E7, levels of IFNκ are elevated in these cells (Fig. 3). In addition to activation of the DDR, the N/Tert-1+HPV16 E6/E7 STOP cells are under replication stress as demonstrated by an attenuation of cellular growth (Fig. 4) and activation of expression of genes involved in replication stress (Fig. 6). These same genes can be activated by E7 when it is over expressed by itself (18). Therefore, there are multiple mechanisms that the virus can use to activate the DDR.

Altogether, the results presented here prompt us to propose the following model: Following infection viral genome copy number increases from 1 to around 50 copies per cell. This increase in copy number would by itself activate the DDR due to torsional stress on the viral DNA. This activation promotes recruitment of homologous recombination factors to the viral genome, a process required for HPV16 replication (47). The activation of the DDR by external DNA damaging agents results in cell cycle arrest to allow detection of damage and repair of the DNA prior to restarting the cell cycle. HPV16 must circumvent this, as it would prevent amplification of the viral genome. Therefore, E6 and E7 manipulate constituents of the DDR to allow cell cycle progression in the presence of an active DDR (this has been previously demonstrated (17, 18, 35, 36)), thus preventing cell cycle arrest. Simultaneously, the actions of E6 and E7 repress innate immune response genes and this not only promotes the viral life cycle via immune evasion, but also abrogates the growth inhibitory activity of innate immune signaling. We show, for the first time, that this targeting of the innate immune response by E6 and E7 is totally synergistic in the context of the entire HPV16 genome; neither protein is able to do this by itself. Therapeutically, targeting only one of the pathways used by E6 or E7 would potentially alleviate the repression of the innate immune response and boost the host immune response to HPV16 infection.

## Materials and Methods

### Generation of HPV16 mutant genomes

Mutations to introduce T584A (E6 STOP), C110T (E7 STOP) or both C110T and T584A (E6E7 STOP) into HPV16 in the peGFP-N1HPV16 plasmid, were carried out by Genscript. Briefly, the target area was released from rest of the plasmid by *Eco*NI and *Not*I restriction digest, base changes introduced and amplified via PCR, which was then religated into the plasmid backbone. Successful mutagenesis was confirmed by sequencing. These mutations resulted in a) an early stop codon in the E6 gene, b) an early stop codon in the E7 gene or c) early stop codon in both E6 and E7 gene (Figure 5).

### Cell culture

Cell lines containing the HPV16 genomes were generated using N/Tert-1 as previously described (30, 31). These cells were cultured alongside parental N/Tert-1 for all comparisons. N/Tert-1 and N/Tert-1+HPV16 cells were grown in K-SFM (Invitrogen) with 1% (v/v) penicillin/streptomycin mixture (Thermo Fisher Scientific) containing 4 μg/mL hygromycin B (Millipore Sigma) at 37°C in a 5% CO2/95% air atmosphere and passaged every 3-4 days. Cells containing WT or mutant HPV16 were grown in the same medium also containing 150 μg/mL G418 (Thermo Fisher Scientific). All cells were routinely checked for mycoplasma contamination.

To measure cell growth, cells were seeded in triplicate onto 10 cm dishes at a density of 3 × 10^5^ cells per dish, and grown to 80% confluency (typically 3 days). Cells were then harvested by trypsinization, stained with trypan blue and viable cells counted. 3 × 10^5^ cells per dish were replated and this was repeated 3 times in total.

For downstream protein and nucleic acid analysis, 1×10^6^ cells were plated onto 100 mm plates, trypsinized and pelleted after 24hrs and washed twice with phosphate buffered saline (PBS).

### RNA and DNA analysis

RNA was isolated using the SV Total RNA Isolation System (Promega) following the manufacturer’s instructions, including the DNAse treatment step. Two micrograms of RNA were reverse transcribed into cDNA using the High Capacity Reverse Transcription Kit (Applied Biosystems). cDNA and relevant primers were added to PowerUp SYBR Green Master Mix (Applied Biosystems) and real-time PCR performed using 7500 Fast Real-Time PCR System. Primer sequences (all 5’ to 3’): HPV16 E2 F atggagactctttgccaacg, HPV16 E2 R tcatatagacataaatccag; HPV16 E6 F ttgaaccgaaaccggttagt, HPV16 E6 R gcataaatcccgaaaagcaa; HPV16 E5 F cacaacattactggcgtgct, HPV16 E5 R acctaaacgcagaggctgct; GAPDH F ggagcgagatccctccaaaat, GAPDH R ggctgttgtcatacttctcatgg; IRF9 F gccctacaaggtgtatcagttg, IRF9 R tgctgtcgctttgatggtact; STAT1 F cagcttgactcaaaattcctgga, STAT1 R tgaagattacgcttgcttttcct; STAT2 F ccagctttactcgcacagc, STAT2 R agccttggaatcatcactccc; IFN-k F gtggcttgagatccttatgggt, IFN-k R cagattttgccaggtgactctt; IFIT1 F agaagcaggcaatcacagaaaa, IFIT1 R ctgaaaccgaccatagtggaaat; MX1 F ggtggtccccagtaatgtgg, MX1 R cgtcaagattccgatggtcct; CHK1 F atatgaagcgtgccgtagact, CHK1 R tgcctatgtctggctctattctg; CHK2 F tctcgggagtcggatgttgag, CHK2 R cctgagtggacactgtctctaa; ATR F ggccaaaggcagttgtattga, ATR R gtgagtaccccaaaaatagcagg; BRCA1 F ttgttacaaatcacccctcaagg, BRCA1 R ccctgatacttttctggatgcc; BRCA2 F acaagcaacccaagtgtcaat, BRCA2 R tgaagctacctccaaaactgtg; RAD50 F tactggagatttccctcctgg, RAD50 R agactgaccttttcaccatgc; OAS1 F tgtccaaggtggtaaagggtg, OAS1 R ccggcgatttaactgatcctg.

DNA was extracted from monolayer grown cells by lysis in HIRT lysis buffer (400 mM NaCl, 10 mM Tris-HCl, and 10 mM EDTA). Cell extracts were incubated with 50 g/ml RNase A, and as well as proteinase K sequentially to remove residual RNA and proteins, followed by phenolchloroform extraction. DNA was resuspended in TE buffer and quantitated by spectrophotometry.Following dilution, 10 ng of DNA and relevant primers were added to PowerUp SYBR Green Master Mix (Applied Biosystems) and real-time PCR performed using 7500 Fast Real-Time PCR System, using sybr green reagent. Primer sequences: HPV16 E2 F 5’ atggagactctttgccaacg-3’ HPV16 E2 R 5’-tcatatagacataaatccag-3’; HPV16 E6 F 5’-ttgaaccgaaaccggttagt-3’ HPV16 E6 R 5’-gcataaatcccgaaaagcaa-3’; HPV16 E5 F 5’-cacaacattactggcgtgct-3’ HPV16 E5 R 5’-acctaaacgcagaggctgct-3’.

### Protein analysis

1×10^6^ cells were lysed in 50 μl NP40 lysis buffer (0.5% Nonidet P-40, 50 mM Tris, pH 7.8, 150 mM NaCl) supplemented with protease inhibitor (Roche Molecular Biochemicals) and phosphatase inhibitor cocktail (Sigma). The cell and lysis buffer mixture was incubated on ice for 20 min, centrifuged for 20 min at 184,000 rfc at 4°C, and supernatant was collected. Protein levels were determined utilizing the Bio-rad protein estimation assay (Bio-rad). Equal amounts of protein were boiled in 2x Laemmli sample buffer (Bio-rad). Samples were then loaded into a Novex 4-12% gradient Tris-glycine gel (Invitrogen), run at 100V for approximately 2 hours, and then transferred onto nitrocellulose membranes (Bio-rad) at 30V overnight using the wet blot method. Membranes were blocked in Odyssey blocking buffer (diluted 1:1 with PBS) at room temperature for 6 h and probed with relevant antibody diluted in Odyssey blocking buffer overnight at 4°C. Membranes were then washed with PBS supplemented with 0.1% Tween (PBS-Tween) before probing with corresponding Odyssey secondary antibody (goat anti-mouse IRdye800cw or goat anti-rabbit IRdye680cw) diluted 1:10000 for 1h at 4°C. Membranes were washed in PBS-Tween before infrared scanning using the Odyssey CLx Li-Cor imaging system. The following antibodies were used for western blot analysis at 1:1000 dilutions in Odyssey blocking buffer, diluted 1:1 with PBS: IFIT1 (D2X9Z), MX1 (D3W7I), and IRF9 (D8G7H) from Cell Signaling Technology. STAT1 (sc-346), STAT2 (sc-1668), pSTAT1 Tyr 701 (sc-135648), pRb (SC-102), p53 (sc-47698) and GAPDH (sc-47724) from Santa Cruz Biotechnology.

### Immunofluorescence

Cells were grown on coverslips to 70% confluence, fixed with formaldehyde and permeabilized with NP40. Cells were incubated with the primary antibody for one hour (phospho-yH2.AX, Cell Signaling Technology 20E3, 1/500), diluted in 10% normal goat serum. Coverslips were washed three times in PBS-Tween (0.1% Tween) and immune complexes were visualized using Alexa 488-or Alexa 595-labeled anti-species specific antibody conjugates (Molecular Probes). Cellular DNA was stained by inclusion of 4’,6-diamidino-2-phenylindole (DAPI, Santa Cruz sc-3598) in the penultimate wash. Microscopy was performed at the VCU Microscopy Facility, supported, in part, by funding from NIH-NCI cancer center grant P30 CA16059. Immunofluorescence was observed using an LSM 700 Laser Scanning Microscope and ZEN 2011 software (Carl Zeiss). Images were assembled in Adobe Photoshop CS 6.0.

### Statistics

Standard error was calculated from at least three independent experiments and significance determined using a student’s t-test.

## Acknowledgements

This work was supported by VCU Philips Institute for Oral Health Research and the National Cancer Institute Designated Massey Cancer Center grant P30 CA016059.

